# Transcription factor action orchestrates the complex expression pattern of *CRABS CLAW*, a gynoecium developmental regulator in Arabidopsis

**DOI:** 10.1101/2021.03.02.433508

**Authors:** Thomas Gross, Annette Becker

## Abstract

The flower of angiosperms is the most complex organ that plants generate and many transcription factors (TFs) are involved to regulate its morphogenesis in a coordinated way. In its center, the gynoecium develops consisting of specialized tissues such as secondary meristems, sites of postgenital fusion, ovules, pollen transmitting tract, all to assure successful sexual reproduction. Gynoecium development requires tight regulation of developmental regulators across time and tissues. However, while we know of several examples how simple on/off regulation of gene expression is achieved in plants, it remains unclear which regulatory processes generate complex expression patterns. Here, we use the gynoecium developmental regulator *CRABS CLAW (CRC)* from Arabidopsis to study regulatory mechanisms contributing to its sophisticated expression pattern. Using a combination of *in silico* promoter analyses, global TF-DNA interaction screens, co-expression and mutant analysis we find that miRNA action, DNA methylation, and chromatin remodeling do not contribute substantially to *CRC* regulation. We show that a plethora of TFs bind to the *CRC* promoter to fine-tune transcript abundance by activation of transcription, linking *CRC* to specific developmental processes but not biotic or abiotic stress. Interestingly, the temporal-spatial aspects of regulation of expression may be under the control of redundantly acting genes and may require higher order complex formation at TF binding sites. We conclude that endogenous regulation of complex expression pattern of Arabidopsis genes requires orchestrated transcription factor action on several conserved promotor sites over almost 4 kb in length.

**Significance statement:** Different to genes that are simply switched on or off, depending on an environmental cue we find that genes directing development in plants often show complex expression pattern dependent on internal factors only. Here, we addressed the question how an complex expression pattern is achieved and use the *CRABS CLAW (CRC)* gene required for gynoecium development as an example. Combining wet lab experiments and *in silico* analysis we find that epigenetic regulation plays only a minor role and that a large number of transcription factors activates the transcription of *CRC*. Single regulators may have a profound effect on *CRC* transcript abundance but less so on the pattern of expression. Complex patterns most likely require the interplay of several transcription factors.

## Introduction

Transcription is a universal process in which DNA is transcribed into mRNA that is exported from the nucleus and translated into protein sequence. Already prokaryotes tightly regulate transcription for a proper timing of cellular development and metabolic processes. However, the prokaryotic way to control of expression is different from eukaryotes, as co-functional genes are often grouped in co-regulated polycistronic operons (reviewed in Riethoven (2010)). In eukaryotes, genes involved in the same process are distributed over the entire genome, such that every gene requires its individual regulatory sequence. Moreover, the promoter regions of eukaryotic genes are longer than those of prokaryotes, include more transcription factor binding sites, accession points for chromatin remodelers, and distal regulatory elements such as enhancers or silencers can be many kilo bases away from the transcription start site (Hernandez-Garcia und Finer 2014).

While the core promoter, which can contain a TATA box or an initiator element, enables the general expression of a gene by recruiting the basic transcriptional machinery (Danino et al. 2015; Porto et al. 2014), the fine tuning of expression is influenced by cis-regulating factors, like enhancers, silencers and insulators (block the action of distant enhancers and mark borders between hetero- and euchromatin), and by trans-regulating factors binding to these elements in the proximal and distal promoter (Zou et al. 2011; Hernandez-Garcia und Finer 2014). Basal transcription factors act as pioneer factors, recruiting additional transcription factors, and opening up DNA binding motifs for specific transcription factors (Thomas und Chiang 2006). The chromatin landscape surrounding the gene directly connects to the ability of transcription factors to bind DNA, such that histone tail modifications influence the accessibility of the chromatin (Lawrence et al. 2016; Deal und Henikoff 2011). Acetylation of histones, e.g. H4K16ac, leads to an opening of chromatin and a higher accessibility of DNA (Lawrence et al. 2016), while the trimethylation of H3K27 leads to condensation of chromatin resulting in reduced transcription, as in the *FLOWERING LOCUS C (FLC)* locus upon vernalization (Bastow et al. 2004).

In addition, DNA methylation suppresses gene transcription. DNA methyltransferases add methyl groups to cytosine residues at three different motifs (CG, CHG, CHH) in plants. If present in promoter regions, DNA methylation usually inhibits enhancer binding and reduces expression (Zhang et al. 2018). The expression of *FLOWERING LOCUS T* is dependent on the distal enhancer *Block C*, while this block is usually demethylated, *FT* expression is inhibited when *Block C* is methylated (Zicola et al. 2019).

Short interfering RNAs (siRNAs) and micro RNAs (miRNAs) are responsible for RNA dependent DNA methylation (RdDM) and post-transcriptional gene silencing (PTGS) (Zhang et al. 2018). While in RdDM siRNA activates *de novo* DNA methylation of complementary DNA regions, PTGS by miRNAs leads to the degradation of complementary mRNAs. Regulation by on miRNAs occurs in multiple genes. For example, members of the HD-ZIP III family are regulated by miRNA 165/166 while miRNA172 binds to the *APETALA2* mRNA (Chen 2004; Miyashima et al. 2011). Regulation of gene expression is thus a combination of diverse regulatory modes, including miRNAs, DNA methylation, histone modifications, and transcription factor activity. However, the contribution of individual aspects of regulation are unknown for most genes. Even more so for genes regulating development of complex organs that require precise spatial and temporal control of expression based mainly on internal signals. And while chromatin modifications, DNA methylation and miRNA binding sites can be measured with precision, transcription factor (TF) binding to specific DNA binding motifs remains elusive. TFs bind to short (6-20 base pair) sequences and those can occur frequently in the genome and be located at random positions. However, only small fractions of the sequences are *bona fide* targets of a particular transcription factor are bound (Lieb et al. 2001), posing major challenges to distinguish the biologically relevant TFBS (TF binding sites) from those that simply match a factor’s binding specificity (Lieb et al. 2001; Moses et al. 2004). Experimental approaches to identify TFBS are ChIP-seq assays (chromatin immunoprecipitation-sequencing) which requires a TF-specific antibody to capture complexes including the TF bound to its target DNA (Kaufmann et al. 2010). However, this is done one TF at the time. To identify all TF’s regulating a single gene’s expression, only few experimental approaches are available, with Yeast One-Hybrid (Y1H) screens where a promoter sequence is used as bait against a TF library being the most extensively used (Yeh et al. 2019). Another option is to identify real TFBS *in silico* by searching for evolutionary conserved sites, as those evolve more slowly than their flanking sequences (Moses et al. 2003). Interestingly, even with abundant gene expression data, TF-binding and TF expression data we lack understanding how changes in TF activity causes changes in target gene expression. Moreover, knowledge on the full set of TFs regulating the complex expression pattern of a developmental regulator in plants based on endogenous cues is unavailable.

The *A. thaliana* protein CRABS CLAW is a member of the YABBY TF family and *crc-8* mutants have a shorter and wider gynoecium with the two carpels apically unfused and they lack nectaries (Bowman und Smyth 1999, Fig. 1). CRC specifies abaxial - adaxial polarity of the carpel, in concert with KANADI proteins and probably antagonistically to members of the HD-ZIP III protein family (Eshed et al. 1999; Reinhart et al. 2013; Tatematsu et al. 2015) and is involved in regulating floral meristem termination. CRC transcriptionally activates carpel target genes regulating nectary formation and gynoecium growth and represses those involved in floral meristem termination (Gross et al., 2018). *CRC’s* expression is strictly limited to the nectaries and the gynoecium (Fig. 1 E-H). In the gynoecium, it commences in stage 6 (stages according to Smyth et al. (1990)) in the gynoecial primordium and forms two distinct domains in the carpels after stages 7-8: n epidermal expression around the circumference of the gynoecium, and an internal expression in four stripes that are close to the developing placenta (Bowman und Smyth 1999; Lee et al. 2005). The epidermal expression of *CRC* is consistent over the complete length of the carpels, but the internal expression forms a basal-apical gradient and ceases in later developmental stages. The epidermal expression is maintained until the mid of stage 12 in the valves, but it decreases earlier in the future replum. Expression in the nectaries starts at their inception and remains stable until after anthesis (Bowman und Smyth 1999). Previous analyzes of the *CRC* promoter by Lee et al. (2005) identified five conserved regions (A-E) sufficient to drive *CRC* expression. Chip-seq data showed that the MADS box transcription factors AG, PI, AP1, and AP3 bind to the *CRC* promoter suggesting their involvement in regulating *CRC* expression (Lee et al. 2005; Gomez-Mena et al. 2005; Ó’Maoiléidigh et al. 2013).

**Figure 1:**
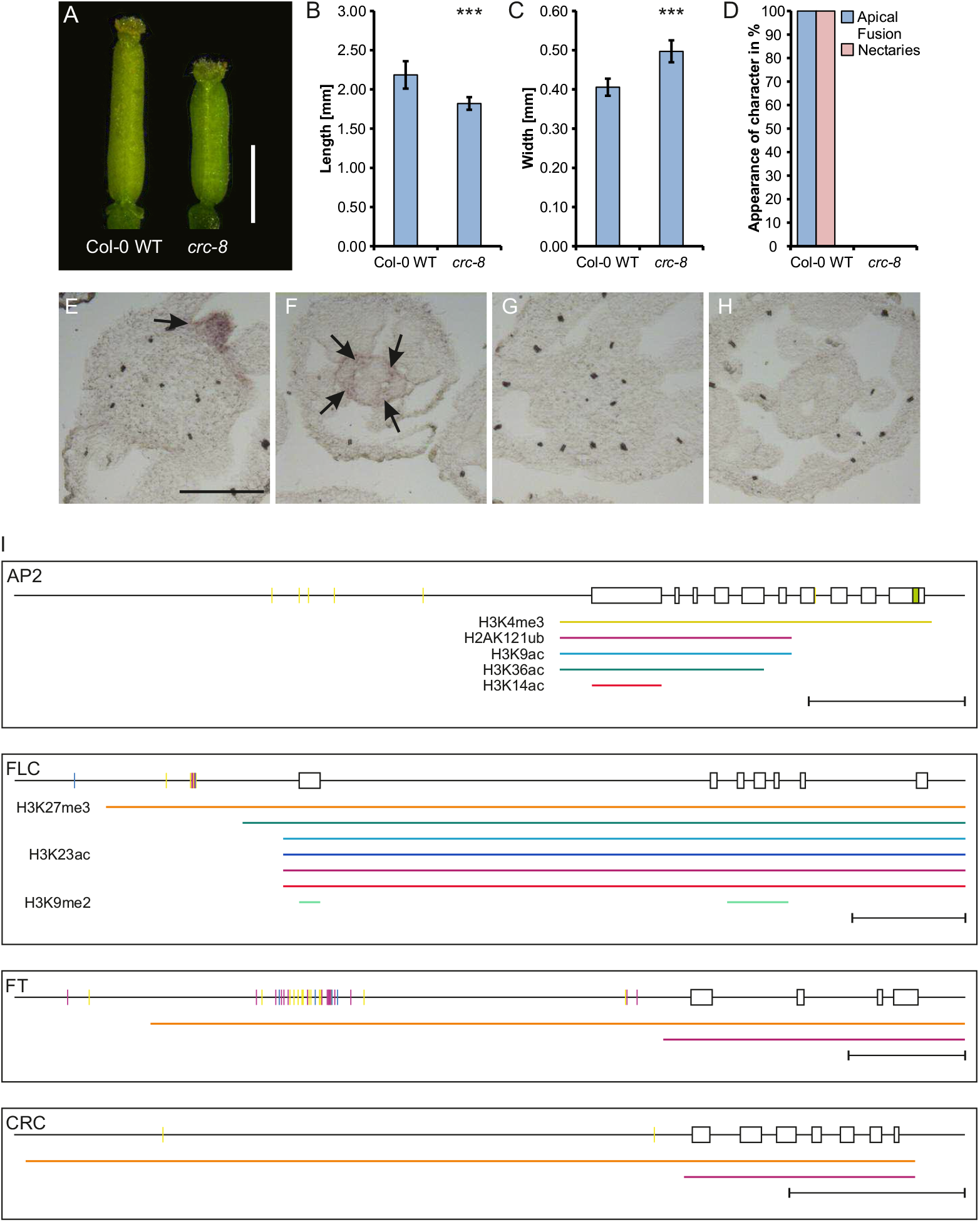
*A. thaliana crc-8* phenotype and summary of gene expression regulation of *AP2, FLC, FT*, and *CRC*. (A) Representative gynoecia of Col-0 wild type and *crc-8* plants. Scale bar represents 1 mm. Statistical analysis of gynoecium length (B), width (C), and a summary of absence or presence of other described *crc-1* phenotypes in *crc-8* (D). Both, length and width comparisons (B, C) are the means with standard deviation. Percent values are shown in (D). Student’s t-test was applied to compare the wild type gynoecia with *crc-8* and significant differences were marked with up to three asterisks (p < 0.001), n=100. (E-H) Spatial analysis of *CRC* expression with RNA *in situ* hybridization. In situ hybridization using a *CRC* antisense probe of *A. thaliana* Col-0 wild type (E and F) and *crc-8* (G and H), showing gynoecia (E and G) and nectaries (F and H). Nectaries and internal *CRC* expression marked with arrows. (I) Summary of gene expression regulation of *AP2, FLC, FT*, and *CRC*. Shown are the promoter regions and the exon/intron structure of the respective gene. DNA methylation is shown in purple (CG), blue (CHG), and yellow lines (CHH). Colored lines under the genomic locus indicate regions of histone modifications identified with PlantPAN: the activating marks H3K4me3 (yellow), H3K9ac (light blue), H3K14ac (red), H3K23ac (dark blue), H3K36ac (dark green), and the repressing marks H2AK121ub (magenta), H3K9me2 (light green), and H3K27me3 (orange). Sorting into activating or repressing marks was performed according to Chen et al. (2015), Mahrez et al. (2016), and Alhamwe et al. (2018). miRNA binding is indicated by a green box in the respective exon. Scale bars represent 1 kB. The ChIP-Seq data used for histone mark identification resulted from only vegetative plant material (seedlings, leaves, roots, and shoot apical meristems/young inflorescence meristems), thus resembling only the state of histone modifications in these tissues.

We were interested in learning more about the regulation of *CRC* expression as an example for a complex expression pattern of a developmental regulator not dependent on external cues. We combine experimental data with *in silico* analysis to identify putative regulators and their roles in *CRC* regulation. We find that TFs involved in several developmental pathways coordinate *CRC* expression via transcriptional activation, such as TFs directing flowering induction, floral organ identity and meristem regulation and that most of them are only partially co-expressed with *CRC*. These regulators bind up to 3 kb upstream of the transcription start site of *CRC*, providing an example showing that complex expression pattern require long promoters

## Results

### *CRC* is not regulated by DNA methylation, chromatin modifications or miRNAs

*CRCs* expression is tightly regulated in a spatial and temporal manner, and specific to carpel and nectary development (Fig 1 E-H). We were first interested in understanding the contribution of the different means of transcriptional regulation of *CRC* expression. In an *in silico* approach, we searched specific databases for DNA methylation sites, histone modifications, and miRNA binding sites in the *CRC*genomic locus (Fig. 1 I) and, in addition, analyzed the genomic loci of *APETALA 2 (AP2), FLOWERING LOCUS T (FT)*, and *FLOWERING LOCUS C (FLC)*. These genes are known to be regulatory by DNA methylation, chromatin modifications and miRNAs, respectively, and serve as controls.

The *AP2* genomic locus show only few DNA methylations between ~2 kB and ~1 kB upstream of the transcription start site (TSS) with five CHH methylations present and an additional CHH methylation at the end of the seventh exon. In contrast to this, multiple CG and CHH DNA methylations were identified ~1 kB upstream of the TSS of *FLC*. In addition, two CHG methylations were present ~1kB and ~2kB upstream of the TSS. The DNA methylation pattern in *FT* is more complex, as methylation marks concentrate on three regions. Approximately 0.5 kB upstream of the TSS are two CG methylation sites and one CHH, 5.5 kB upstream with only few DNA methylations (CG and CHH) in the *FT* promoter and in a highly methylated stretch of ~1 kB between 2.7-3.7 kB upstream of the TSS, including many CG and CHH methylations but also few CHG methylations. The *CRC* genomic locus shows two sites of CHH DNA methylations (~0.3 kB and ~ 3 kB upstream of the TSS), suggesting little influence of DNA methylation on its gene expression. Activating and repressive histone marks were found in the genomic loci of all four genes based on ChIP-Seq data from vegetative plant tissues. Both, the genomic loci of *AP2* and *FLC* showed the highest number of histone marks and included activating and repressing marks covering most of promoter regions and the coding sequences. In contrast, *CRC* and *FT* genomic loci show only two repressive marks, with H3K27me3 covering most of the genomic locus, including the promoter and H2AK121ub covering only the transcribed region. MiRNA binding sites were identified only in the last exon of *AP2*.

In summary, our *in silico* analysis corroborates that *AP2* is regulated by all analyzed means of expression regulation. *FLC* shows regulation by activating and repressive histone marks as well as CHH, CG, and CHG DNA methylation. *FT* has only few types of repressive histone marks present and is regulated by extensive CHH, CG, and CHG methylation. In contrast, *CRC* is regulated independently of miRNAs and DNA methylation. Similar to FT, is shows only two types of repressive histone marks in vegetative stages of development.

### Diverse transcription factors bind to the *CRC* promoter

Because we have shown that the genomic locus of CRC is not affected DNA methylation, only mildly by histone modifications, and that miRNA cleavage of its transcripts also play no role for gene expression regulation (Fig. 1 I) we hypothesized that *CRC* is regulated mainly by transcription factors. To identify the direct upstream regulators of *CRC*, a Yeast-1-hybrid (Y1H) screen of the *CRC* promoter was performed in which the full-length promoter (3.8 kb) and 12 smaller promoter fragments (Suppl. Fig. 1) were transferred as baits into yeast. Four bait strains (*proCRC A, proCRC F2*, and *proCRC F5*) were discarded because they showed autoactivation with resistance to 1000 ng/ml AbA. The remaining ten strains were transformed with the three different libraries of prey TFs (Suppl. Table 3 and Mitsuda et al. (2010)) and grown on selective SD-Leu or SD-Trp medium. 140 proteins binding to the *CRC* promoter fragments in yeast were identified (Suppl. Table 4). We used PlnTFDB (Pérez-Rodríguez et al. 2010) and TAIR database to these TFs to protein family. To identify those that bind sequence specifically, we searched PlantPAN 3.0 (PP, Chow et al. 2019) for the presence of their experimentally verified DNA binding motifs in the *CRC* promoter (Suppl. Table 4).

34 % (48 proteins) of the 140 proteins are present in the PP database and have a DNA binding motif match within the *CRC* promoter sequence (Fig. 2 A). The remaining 92 prey proteins could be separated into two categories: 1) 71 proteins (51 %) were not included in PP, because their DNA binding motif is unknown, 2) 21 proteins (15 %) were included in PP but their binding motifs do not match to the CRC promoter. The 21 proteins from the second category may have additional binding motifs not yet identified, or they have indeed no binding site in *proCRC* and can be seen as false positives and were thus excluded from further analyses. All genes encoding proteins identified in this Y1H screen are expressed during gynoecium development (Schmid et al. 2005; Klepikova et al. 2016) and resemble multiple protein families (Suppl. Figure 2). The proteins binding to the *CRC* promoter in yeast include well-known carpel developmental regulators like HALF FILLED (HAF), FRUITFUL (FUL), ETTIN (ETT) and ARF8. But also genes so far not known to act in the gynoecium, such as REVEILLE 4 (RVE4), required for circadian rhythm maintenance and response to heat shock (Li et al., 2019; Gray et al., 2017), or WRKY41, involved in regulation anthocyanin biosynthesis (Duan et al., 2018) (Fig. 2A). These results suggest that *CRC* expression regulation relies on several, so far seemingly unrelated developmental pathways

**Figure 2:**
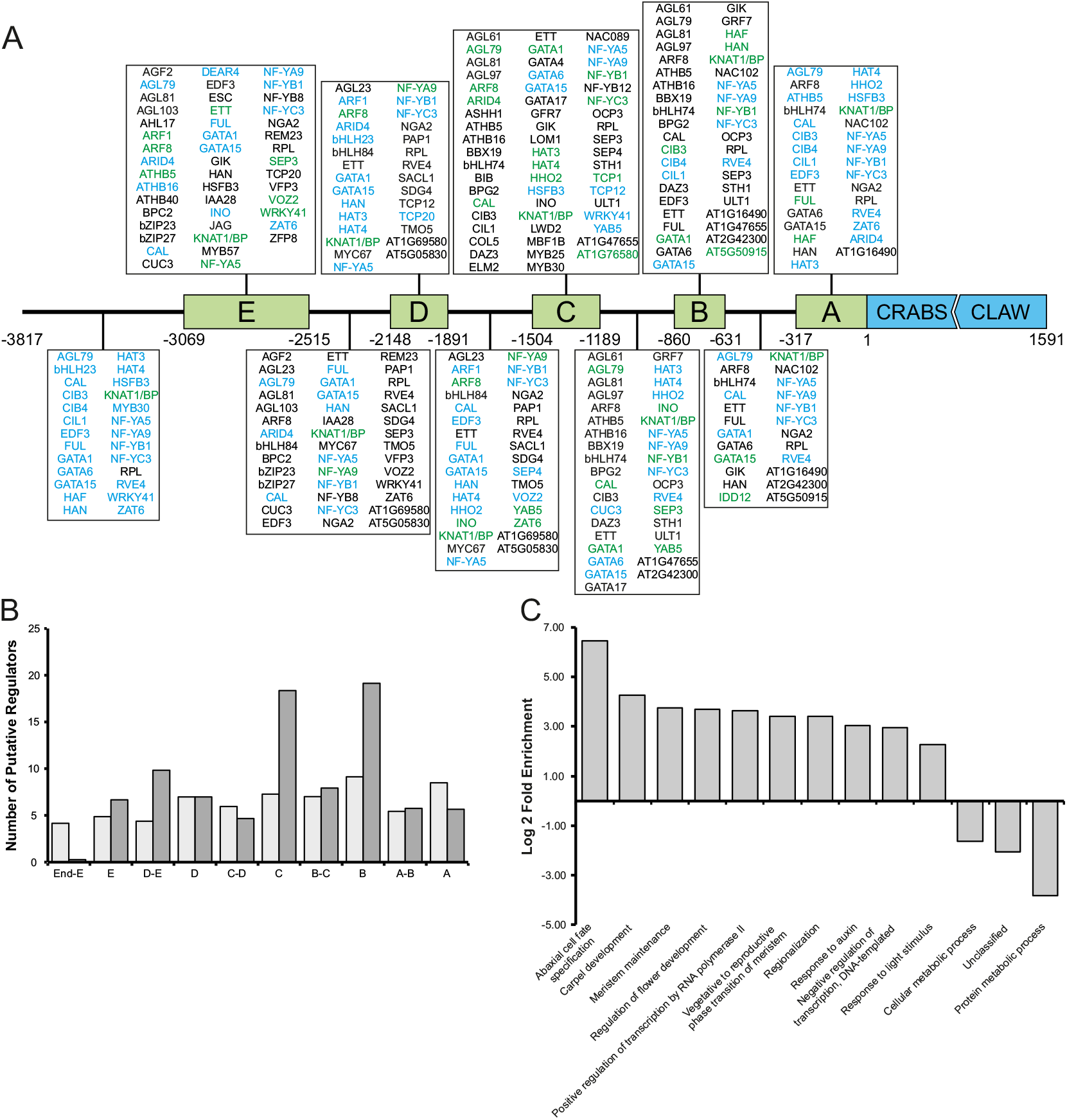
Analysis of transcription factors binding to the *CRC* promoter identified by Yeast One-Hybrid analysis. A) Spatial distribution of transcription factor binding sites summarizing the Y1H screen of *proCRC* with transcription factor bait libraries using the fragments indicated in Supplemental Fig. 1 as prey. Shown are only transcription factors with a known motif in PlantPan. Binding sites proteins in black were identified by Y1H, those in blue by PlantPAN *in silico* prediction, and those in green indicate positional overlap of Y1H and PlantPAN data. B) Quantitative analysis of putative *CRC* regulators distribution across the different fragments of *proCRC*. The number of regulators identified by in silico prediction with PlantPAN per 100 bp is shown in light grey and the number of transcription factor binding sites identified by the Y1H screen per 100 bp are shown in dark grey. C) GO enrichment analysis, categorizing the putative *CRC* regulators in different functional groups. Shown is the log 2-fold enrichment of significantly overrepresented GO terms.

### Relevant promoter fragments are enriched in TFBS and *CRC* regulators are functionally related

We were then interested to see if the binding sites of the direct regulators are located in the regions conserved between Brassicaceae (Lee et al., 2005). We plotted the number of TF binding sites per 100 bp in one region identified via *in silico* prediction (light grey) and identified in the Y1H screen (dark grey) (Fig. 2B). On average, 6.37 and 8.52 binding sites per 100 bp were identified, fragments B (9.13 and 19.13, respectively) and C (7.28 and 18.35, respectively) show most binding sites, with an additional maxima in A for the in silico identified binding sites (8.49 and 5.66, respectively). Overall, the fragments including the fewest putative binding sites are all between the conserved blocks identified by Lee et al., (2005a) corroborating their results from promoter shading and promoter-GUS assay analysis.

We were then interested to learn about the function of the *CRC* regulators. Thus, the 119 putative *CRC* regulators (those identified with Y1H and known DNA binding motifs in *proCRC* plus those of category 1) were divided into functional groups based on gene ontology terms (Figure 2C). Ten GO terms were overrepresented among the candidate genes (abaxial cell fate specification, carpel development, meristem maintenance, regulation of flower development, positive regulation of transcription by RNA polymerase II, vegetative to reproductive phase transition of meristem, regionalization, response to auxin, negative regulation of transcription (DNA-templated), and response to light stimulus), while three GO terms were underrepresented (cellular metabolic process, unclassified, and protein metabolic process). Most enriched terms are closely related to known functions of *CRC*, especially the terms abaxial cell fate specification (GO:0010158) and carpel development (GO:0048440) are highly enriched, with a 6.47 log 2-fold enrichment and 4.28 log 2-fold enrichment respectively. Only the weakly enriched (2.26- log 2-fold) response to light stimulus (GO:0009416) is not directly related to *CRC* functions but might be connected to light induced flowering, through genes like flowering time regulators *LIGHT-REGULATED WD2 (LWD2)* and *VASCULAR PLANT ONE ZINC FINGER PROTEIN 2 (VOZ2)*..

### *CRC* expression is activated by diverse developmental regulators

We were then interested in how the *CRC* regulators influence the pattern and strength of CRC quantitatively *in planta. CRC* expression in *hbb* and *cauliflower (cal)* was visualized using mRNA *in situ* hybridization and showed no differences between to the wild type expression (Suppl. Fig. 4).. Interestingly, no *CRC* expression was found in the newly characterized *crc-8* (Fig. 1, Suppl. Fig. 4), suggesting auto-activation and/or –maintenance of *CRC* transcription. Further, we chose four mutants (*agf2; Arabidopsis thaliana homeobox 16, athb16; bbx19; indeterminate domain 12, idd12; inner no outer, ino*) based on their defects in flower development or phytohormone signaling (Suppl. Table 1) for GUS staining assays. The localization of *CRC* expression in all four mutants was similar to the wild type suggesting that the loss of function of those single regulators has no effect on the spatiotemporal expression of *CRC* (Suppl. Figure 4).

We also quantified changes in *CRC* expression in 20 homozygous regulator mutants via qRT-PCR (Fig. 3A). *CRC* transcript levels in buds are significantly reduced in 14 mutants: *agf2, bbx19, cal, ettin (ett), fruitful (ful), half-filled bee1 bee 3* triple mutant *(hbb), hat4, idd12, jagged (jag), nf-ya9, ngatha2 (nga2), reveille4 (rve4), ultrapetala1 (ult1)*, and *yabby5 (yab5)* (Figure 3 A). Of those, *CRC* expression was decreased by only ~25 % in *hat4* (0.78 +- 0.11), *idd12* (0.78 +- 0.16), *rve4* (0.74 +-0.15), and *ult1* (0.74 +- 0.15) when compared to wild type expression. The other regulator mutants showed *CRC* expression reduction between25 % - 50 %, for example in *ett* buds, *CRC* expression was only half as strong as in wild types buds (0.46 +- 0.12). These findings indicate that these transcription factors activate expression of *CRC* in the gynoecium and/or nectaries.

**Figure 3:**
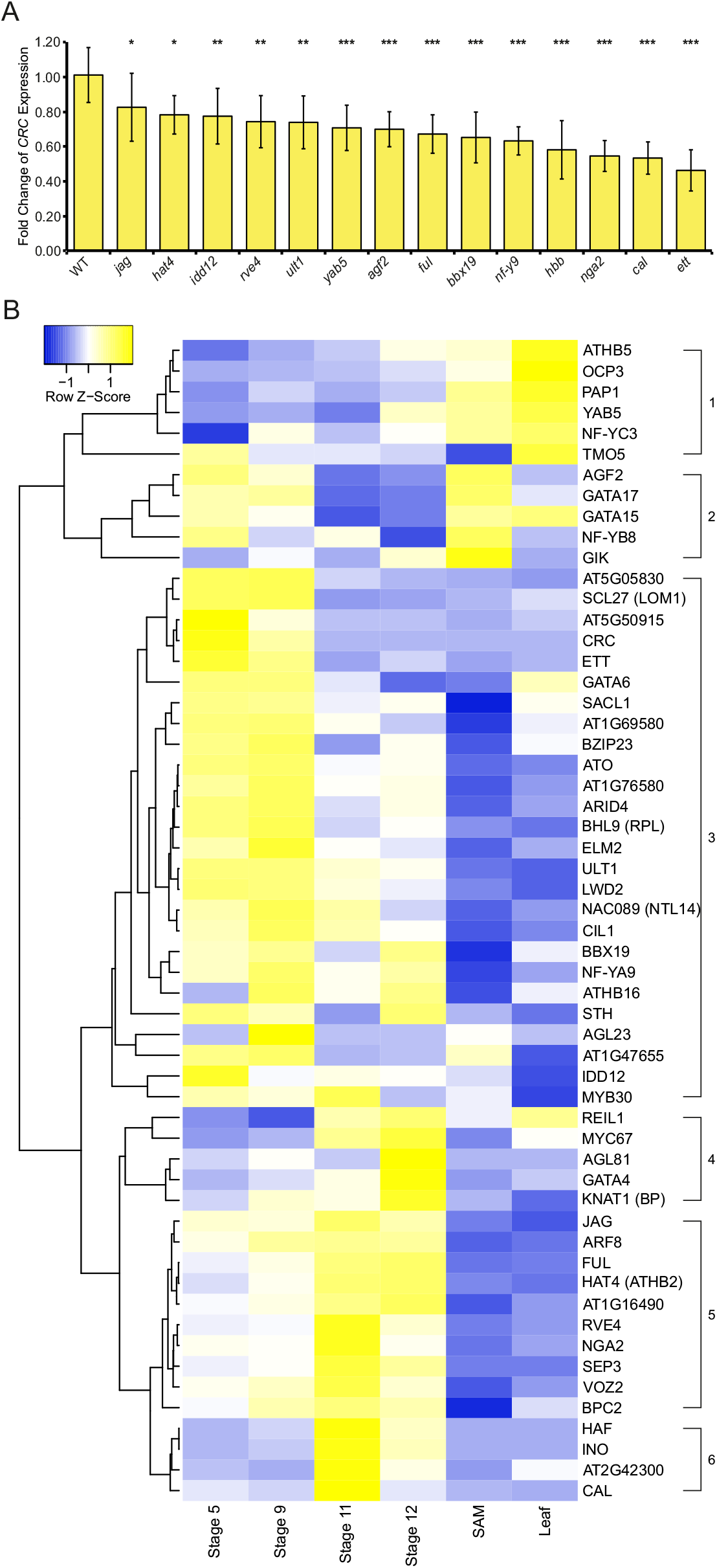
Quantitative effect of regulator mutants on *CRC* expression and digital gene expression analysis of putative *CRC* regulators. A: *CRC* expression in candidate mutant lines in relation to *CRC* expression of Col-0 wild type buds given as mean values of the fold change of *CRC* expression, error bars indicate standard deviation. B: Heatmap of *CRC* and co-expressed putative regulators during four carpel developmental stages (Kivivirta et al., 2020), shoot apical meristem (SAM) and leaves (Klepikova et al., 2016). The rows are not correlated to each other. Color intensity represents z-score.

*Auxin response factor 8 (arf8), athb16, cib1-like protein 1 (cil1), ino*, and *target of monopteros 5 (tmo5)* mutants showed *CRC* transcript abundance similar to the wild type (Suppl. Fig. 5). This suggests that six proteins of 20 bind to the *CRC* promoter in yeast and have predicted binding sites in this promoter but do not contribute to *CRC* expression regulation, while 14 proteins activate *CRC* expression. Among the activators are at least four (AGF2, BBX19, CAL, HAF) that, as single genes, have no influence on the pattern of expression but on transcript abundance.

### Regulators of *CRC* are partially co-expressed during flower development

As several methods for TF target prediction use co-expression of the TF and its target genes (e.g. Margolin et al., 2006) we were interested to learn how useful this method is for identifying regulators of complex expression patterns. We thus carried out digital gene expression analysis of the candidate regulators to discriminate genes with expression patterns similar to *CRC* from those with complementary expression patterns. 7577 genes were co-expressed with *CRC* based on Pearson correlation (correlation coefficient between 0.8 – 1 and −0.8 – −1), and among those, 5167 were positively and 2410 negatively correlated with *CRC* expression. 555 of the co-expressed genes are TFs, including 381 positively and 174 negatively correlated genes. 32 co-expressed genes encoded transcription factors binding to the *CRC* promoter shown in the Y1H experiments (Fig. 2). To these 32 co-expressed genes we added the candidate regulators chosen for mutant analysis, if they were not already present (Fig. 3 B). These co-expressed genes assemble into six groups: group 1 includes six genes that are mainly expressed in leaves. Group 2 genes are highly expressed in early carpel stages and SAM with five members, two of them are the closely related GATA transcription factors GATA15 and GATA17. Genes expressed mainly in early carpel developmental are members of group 3, including CRC and most other genes encoding for proteins shown to interact with proCRC in the Y1H screen including well known genes like *ETT, BHL9 (RPL)* and *ULT1*. Group 4 members are mainly expressed in the latest stage of carpel development and include for example *KNAT1*. Group 5 members are most strongly expressed in the carpel at stage 11 of flower development and include genes like *JAG, ARF8, FUL, NGA2*, and *SEP3*. Group six includes only four genes with almost exclusive expression in stage 11 including *HAF, INO*, and *CAL*. The inclusion of additional RNAseq data from inflorescences, young flowers, and mature flowers further subdivided the six categories into nine (Suppl. Fig 3)

In summary, we find that the largest group of proteins interacting with the *CRC* promoter shows an expression pattern very similar to that of *CRC* indicating a role in activation or maintenance of *CRC’s* expression. Other groups (1 and 2) show partially or fully complementary expression pattern suggestive of a function in negative regulation of *CRC*.

## Discussion

### A complex interplay of transcription factors regulates *CRC* expression

*CRC* is expressed in a complex spatial and temporal manner, and as *CRC* expression is not regulated by DNA methylation, miRNAs or repressive histone marks (Fig. 1 I), its spatial and temporal expression regulation relies mainly on TF networks (Fig. 4A).

**Figure 4:**
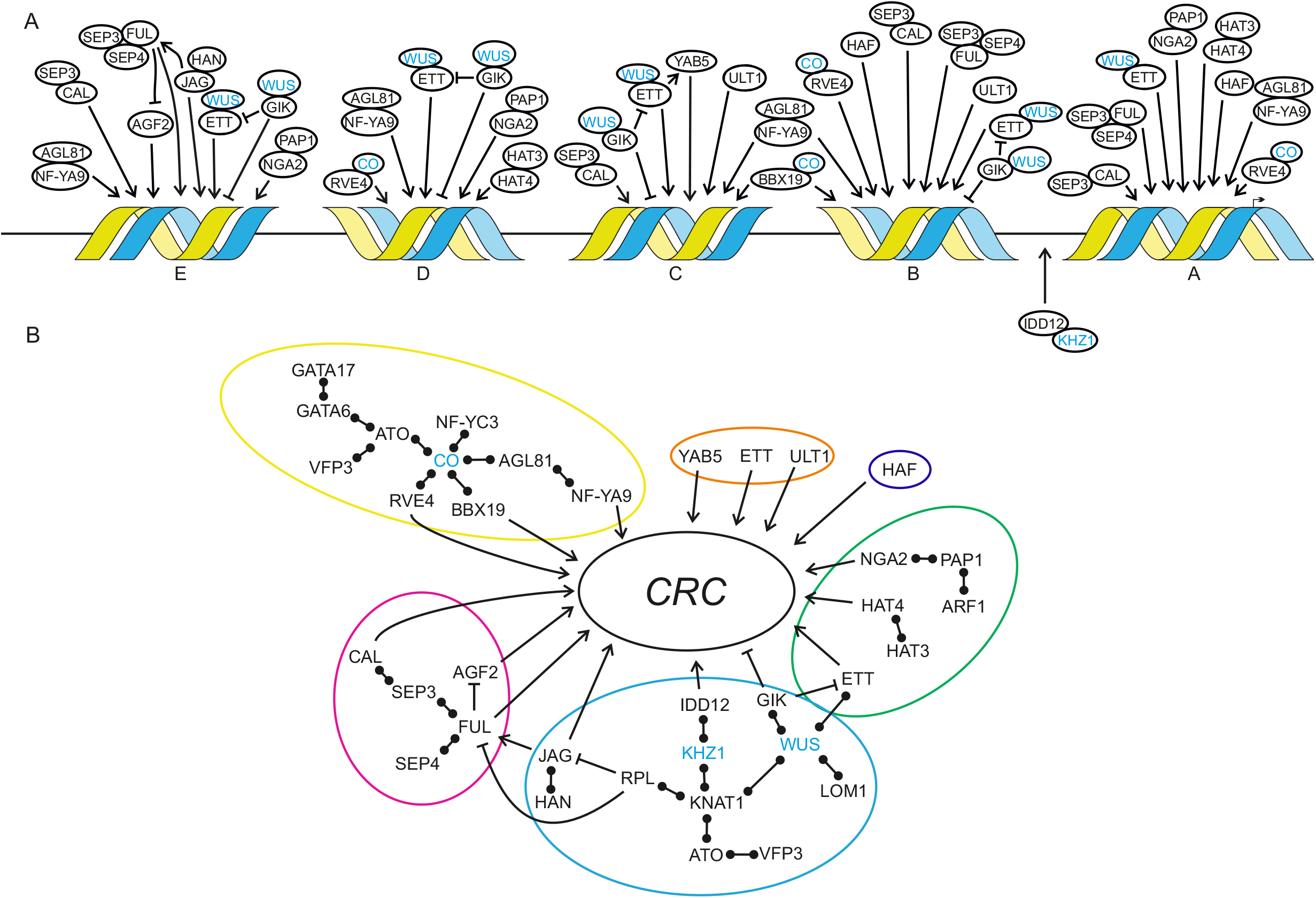
Networks regulating *CRC* expression. A) Localization and binding of *CRC* regulators based on Y1H and qRT-PCR results and Ng et al. (2009). Regulators were assigned to the conserved regions of *proCRC* (A-E) accordingly to the Y1H screen and *in silico* prediction analyses. Proteins without an arrow were not tested via qRT-PCR but interact with proteins identified in this study based on BioGRID searches (Oughtred et al. 2019). Protein-protein-interactions are symbolized by overlapping circles and proteins in blue were not present in the Y1H results, but link identified regulators to each other. Protein interactions are shown by overlapping circles, transcriptional activation and repression are symbolized by pointed or blunt-end arrows, respectively. B) Contribution of flower-related processes to *CRC* expression. Colored circles indicate co-functional networks with flowering induction in yellow, ab-/adaxial regulation in orange, carpel structures in dark blue, auxin response in green, meristem regulation in light blue, and flower development in purple. Protein interactions are shown by bars with two circles, transcriptional activation and repression are symbolized by pointed or blunt-end arrows, respectively.

*CRC* is integrated in different regulatory networks necessary for flower development, like the termination of the floral meristem or adaxial/abaxial polarity of carpel development (Bowman und Smyth 1999; Prunet et al. 2008; Sun und Ito 2015) (Fig. 4B). The members of gene regulatory networks are often co-expressed and co-expression analysis may be used to carefully predict the function of a gene, as co-expressed genes are not necessarily co-functional (Usadel et al. 2009).

Here, we combine data from the co-expression analysis and Y1H screen (Fig. 2 and 3) with data derived from literature to maximize the likelihood to find “real” regulators of *CRC*.

Among the putative repressors of *CRC* in leaves and the SAM are ATHB5 and ATHB16, which can heterodimerize (Johannesson et al. 2001; Wang et al. 2003), and they are both strongly expressed in the SAM and leaves. ATHB5 is involved in hypocotyl development and represses other transcription factors like *BODENLOS* (Smet et al. 2013). ATHB5 and ATHB16 proteins bind to regions C and B of the *CRC* promoter (Fig. 4A), and *CRC* expression in *athb16* buds remains unchanged (Suppl Fig. 5) suggesting that they may repress *CRC* expression as heterodimer in vegetative tissues. *NUCLEAR FACTOR-Y C 3 (NF-YC3)* is also strongly expressed in vegetative tissues and involved in the regulation of photomorphogenic growth by recruiting histone deacetylases to the chromatin regions of its target genes (Tang et al. 2017). The stable repression of *CRC* in the vegetative phase of *A. thaliana* might be thus achieved by combinatorial action of transcriptional repressor activities and repression by chromatin deacetylation (Roth et al. 2001; Kouzarides 2007).

Among the regulators co-expressed with *CRC* in the gynoecium and a strong expression in the SAM is AGF2, which activates *CRC* when FUL expression is low (Fig. 3A and B). Also GIANT KILLER, an AT-hook type DNA binding protein, which is known to counteract *ULT1* activity, (Ng et al. 2009) shows this peculiar expression pattern. GIK expression is directly activated by AG and it is known to regulate genes involved in carpel development like *ETT*, by adding the repressing histone marks H3K9me2 to the *ETT* promoter (Ng et al. 2009). GIK binds *proCRC* fragments C, D, and E may repress *CRC* expression late stages in gynoecium development in a way similar to *ETT*.

ULT1, a SAND and trithorax domains containing transcriptional regulator (Bottomley et al. 2001; Carles und Fletcher 2009)), activates *CRC* expression in flowers (Fig. 2, Fig. 3A). It mediates the removal of repressive histone H3 lysine methylation marks (H3K27me3) or hinders their new positioning and to activate the expression of its target genes, such as *AG* (Carles und Fletcher, 2009). As ULT1 binds to the *CRC* promoter regions B to C, it may mediate the removal of repressive histone marks to the *CRC* genomic locus early in gynoecium development, in a way similar to what was shown for *AG* (Carles and Fletcher, 2009). *ULT1* also indirectly activates *CRC* by activation of *AG*, which, in turn activates *CRC* expression (Bowman und Smyth 1999; Lee et al. 2005; Ó’Maoiléidigh et al. 2013). Interestingly, *CRC* and *ULT1* act redundantly to terminate the floral meristem (Prunet et al. 2008) suggesting that *ULT1* and *CRC* act on the same targets while *CRC* itself is a target of *ULT*.

ETT binds to the promoter fragments A-E in yeast (Fig. 2A) and its loss of function results in the strongest decrease of *CRC* expression when compared to all other genes tested (Fig. 3A), suggesting that ETT is an important activator of *CRC* transcription. In leaves, ETT activates the expression of the YABBY genes *FIL* and *YAB3* which act in combination with KAN genes in specifying abaxial polarity (Garcia et al. 2006). Because in carpels it is *CRC*, which is involved in abaxial polarity specification, *ETT* may target YABBY genes in a more general way. Interestingly, *ETT* is only weakly co-expressed with *CRC* and provides an example of an important transcription factor not strongly co-expressed with its target.

Interestingly, several genes activating *CRC* expression are only weakly co-expressed with *CRC*. Among those are *FUL, HAF, NGA2*, and *JAG*, which are all most strongly expressed in later stages (Fig. 3B) of carpel development but strongly activate *CRC* expression (Fig. 3A) and bind to *proCRC* (Fig. 2A). The bHLH protein HALF FILLED is necessary for transmitting tract development to enable the pollen tubes growth for ovule fertilization (Crawford und Yanofsky 2011; Crawford et al. 2007). FUL acts antagonistically to RPL and together they determine valve identity and are necessary for the elongation of the developing fruit (Gu et al. 1998; Ferrandiz et al. 2000). A ChIP-SEQ analysis of FUL targets did not identify *CRC* (Bemer et al. 2017), but only gynoecia and fruits after stage 12 were part of this analysis suggesting that FUL may be absent from the *CRC* promoter in late stages of carpel development. NGA2 binds to regions E and A of *proCRC*, it participates in the formation of style and stigma and is involved in longitudinal growth of the gynoecium ((Ballester et al. 2015; Lee et al. 2015) Alvarez et al. 2009; Trigueros et al. 2009). As *crc* gynoecia are typically shorter than those of the wild type, NGA2 might act via activating *CRC* to control this longitudinal growth.

Also, JAGGED (JAG) activates *CRC* expression while it is only weakly expressed during early gynoecium development and it is genetically interacting with several co-expressed proteins. This group of genes may act in a concentration-dependent regulation such that FUL, HAF, NGA2, and JAG activate *CRC* at low, and repress *CRC* at high protein concentration.

As expected, MADS-box proteins are involved in *CRC* regulation: SEP3 and SEP4 physically interact with AG, AP1, and PI, which are known to regulate *CRC* (Immink et al. 2009;Bowman und Smyth 1999; Ó’Maoiléidigh et al. 2013). Thus, their binding sites can serve as hubs for MADS regulation. Also CAL binds to *proCRC* in its C region, but is not known to have roles late in flower development. It acts redundantly with *AP1* to orchestrate the transition from inflorescence meristem to floral meristem and is expressed mainly in the floral meristem, sepals and petals (Alvarez-Buylla et al., 2006). However, *CAL* is also necessary to activate other flower developmental genes and may be involved in the initiation of *CRC* expression at low dosage (Fig. 3A). In addition, *CAL* may have a repressive function on *CRC* late in gynoecium development when it shows a peak of expression at stage 11 (Fig. 3B) suggesting a dosage-dependent action on *proCRC*.

*CRC* expression is regulated by members of different developmental networks connected by protein interactions, which are all involved in reproductive development (Fig. 4B): the genetic network regulating floral induction by light with CO as central regulator acts on *proCRC* directly via RVE4, BBX19, and NF-YA9. The floral meristem regulation network acts on *proCRC* via JAG, IDD12 and GIK, and the floral organ identity network via CAL, FUL, and AGF2. Our data show that CRC is not only directing auxin synthesis and readout by repressing *TRN2*, a modulator of auxin homeostasis, and by regulating *YUC4*, an auxin biosynthesis gene (Yamaguchi et al., 2018), but it is also regulated by the auxin response factor ETT, HAT4, and NGA2, the latter being involved in auxin signaling (Fig. 4B). Also, adaxial/abaxial polarity regulators such as ETT, ULT1, and YAB5 activate *CRC* expression, as well as HAF, which a gynoecium morphogenesis regulator. Proteins described as members of the gynoecium morphogenesis network (Kivivirta et al., 2020) are also participating in other networks (Fig. 4B), such as SEP3, FUL, ETT, NGA2, KNAT1/BP, or RPL regulate *CRC* expression, suggesting that the *CRC* promoter receives signals from several interconnected developmental GRNs allowing precise timing, spatial distribution, and control of transcript abundance for proper *CRC* expression.

### Regulation of complex expression patterns

The regulation of a single gene’s expression at the level of timing, distribution, and abundance of transcripts in a comprehensive way is addressed in surprisingly few studies. For genes responding to external cues like heat stress, interaction between auto- and cross regulation of TFs, epigenetic and post-transcriptional regulation is combined to acquire thermotolerance and long-term adaptation to heat stress (reviewed in Ohama et al., 2017). But these genes are expressed rather uniformly in the plant or at the site of induction. In contrast, developmental regulators respond to internal cues, such as regulation by other transcription factors and for some of them, one being *CRC*, these internal cues of assumedly several pathways are summed up and produce intricate mRNA patterns in space and time. Many examples of these developmental gene regulatory networks (GRNs) are known, e.g. for flower development (Thomson and Wellmer, 2019), or leaf development (Du et al., 2018) or the precise spatial patterning of lignification allowing for fruit dehiscence (Ballester and Ferrandiz, 2017). However, most of these studies used co-expression analyses or reconstructed GRNs by assembly of single/few genes’ genetic interactions. In contrast, our study provides evidence independent of genetic interaction or co-expression studies on the complex regulation of complex expression patterns of a key developmental regulator. Our work indicates that the combinatorial action of TFs is important for patterning of expression, but less so for the regulation of transcript abundance. This leads to the question if TF’s transcript abundance is of lesser importance than the spatial distribution of transcripts. Many genes are thought to be differentially expressed with a log2 fold change difference (e.g. Huang et al., 2019, Hackett et al., 2020) between treatments or developmental stages. However, one may ask how relevant this threshold is for developmental regulators, which may act at low abundance. However, other TFs like WUS or PLE are known for their dosage dependence as high transcript/protein abundances result in binding of low affinity binding sites and low protein abundances results in high affinity binding site usage (reviewed in Hofhuis and Heidstra (2018)).

GRN members generally seem to be co-expressed and this knowledge is used to link unknown genetic connection to GRNs *in silico*. However, our work shows that this approach may be used only with utmost care as most RNAseq data (e.g. Klepikova et al., 2016; Chen et al., 2018) lack the precision for taking tissue- or cell-type-specific expression into account or are not developmental stage-specific. Further, many developmental regulators act in a tissue context-dependent way, together with different protein interaction partners, and on several target genes. For example, the *CRC* activator ETT interacts with TEC1 to repress the growth of side shoot structures in an auxin dependent manner (Simonini et al., 2017) and interacts with ABERRANT TESTA SHAPE (ATS or KAN4) to define boundaries between integument primordia of ovules (Kelley et al., 2012). Within the tissues selected for the co-expression analysis (Fig. 3B), *ETT* and *CRC* are co-expressed, but if inflorescence axis, hypocotyl, and ovules would have been added, together with the ovule-free carpel datasets from Kivivirta et al. (2021), co-expression between the two genes would be difficult to find, as *ETT* shows expression in those tissues (Klepikova et al., 2016) but *CRC* does not (Suppl. Fig. 3). The same is true for *ATHB16*, a gene that is co-expressed with *CRC* in our datasets but does not regulate *CRC* (Suppl. Fig. 6) and would most likely be a false positive member of the *CRC* containing GRN. Conversely, *CAL* is hardly expressed in the gynoecium but activates *CRC* expression (Fig. 3A). This shows that co-expression and co-regulation are not necessarily linked and that the assembly of GRNs based on co-expression networks requires extensive experimental verification.

While we tried to obtain a comprehensive picture of experimentally verified, direct regulators of *CRC* expression, the Y1H approach identifies only transcription factors acting as monomers or homodimers. To our knowledge, there is no published information on the ratio of TFs forming homo- vs. heterodimers in plants. However, a compilation by Kivivirta et al. (2020) shows that among the known interactions of carpel development regulators, only 25 can homodimerize but 56 cannot and that dimeric interactions change over time. If TFs can bind to DNA only in a specific combination with another TF, Y1H will not identify this interaction. Once a homodimer is identified, higher levels of complexity can be analyzed by a combination of protein interaction data resulting from other methods, such as Yeast Two-Hybrid (Y2H) or CrY2H-seq (Trigg et al., 2017) (Fig. 4).

Our data further show that transcription activating TFs, such as ETT bind up to 3 kb upstream of the transcription start site (Fig. 2, Suppl. Fig. 1, Lee et al., 2005). These experimental findings do not corroborate *in silico* analyses (e.g. Yu et al., 2016) showing that the majority (~80%) of TFBS of the majority of promoters are between −1000 and +200 of the transcription start site. However, genes with complex expression patterns may be also the exception to the general positional preferences with more extended promoters. In addition, many methods used to identify transcription factor DNA binding motifs take only monomers or homodimers into account and thus show often palindromic binding motifs. However, the formation of heterodimers can influence directly the DNA binding motifs of the two dimerizing proteins (Jolma et al. 2015, Inukai et al. 2017), thus increasing the difficulty of *in silico* binding site predictions.

In summary we can state that the comprehensive analysis of factors regulating complex transcription factors is, at the current state of wet lab and *in silico* methods, challenging. Directed manipulation of developmental regulator’s expression pattern for yield improvement is thus difficult to achieve and requires extensive research.

## Materials and Methods

### Plant Material and Plant Growth

All plants were grown on a soil-perlite mixture under standard long day conditions. For the crosses with the GUS reporter line, SALK lines of various transcription factors were used (Suppl. Table 1). For RNA *in situ* hybridizations, the SALK line (SALK_007052C, in Col-0) with a T-DNA insertion in the sixth exon of *CRC* (henceforth *crc-8*), the *half filled, bee1, bee3* triple mutant (*hbb*) (a kind gift of Birgit Poppenberger and Martin Yanofsky), and the *cal* mutant (a kind gift of Daniel Schubert) were used. For a detailed description of *crc-8*, 100 randomly picked flowers at stage 14 (Smyth 1993) *A. thaliana* Col-0 wild type plants and *crc-8* plants, respectively, were manually dissected under a Leica M165C stereoscope (Leica Microsystems GmbH, Wetzlar, Germany) analysed (Fig. 1 A-H).

### Y1H assay

The *CRC* promoter (*proCRC*), as described by Bowman und Smyth (1999) and Lee et al. (2005), was amplified as a 3.8 kB fragment from genomic DNA of *A. thaliana* Ler-0. Additionally, the promoter was divided into seven fragments (*proCRC F1 – F7*) and into the five conserved regions (*proCRC A – E*) (Suppl. Fig 1) that were identified by Lee et al. (2005) were PCR amplified (for primers see Suppl. Table 2), digested with HindIII and KpnI, and cloned into the equally digested bait DNA vector pAbAi (Takara Clontech, Saint-Germain-en-Laye, France). The yeast strain *S. cerevisiae* Y1HG (Takara Clontech) was used for all Y1H analyses. The yeast transformation and autoactivation test was performed as described in Gross et al. (2018). The lowest Aureobasidin A (AbA) concentration that was sufficient to suppress yeast growth was used for the following screens. AbA sensitive strains were then transformed with the three prey libraries (for compositions of the three libraries see Suppl. Table 3 and Mitsuda et al. 2010). Prey plasmids from colonies >2 mm diameter were isolated and sequenced.

### Construction of *proCRC:GUS* reportersystem and GUS assays

As *proCRC* exhibits an internal BsaI recognition site, site-directed mutagenesis (Hemsley et al. 1989) of *proCRC* was performed to remove the BsaI recognition site (primers are listed in Suppl. Table 2) for the later integration of *proCRC* into the Greengate system (Lampropoulos et al. 2013). After integration, the construct *proCRC:N-Dummy:GUS:C-Dummy:TerRBCS;pMAS:Basta:TerMAS* in the plant transformation vector pGGZ003 was assembled as described in Lampropoulos et al. (2013). The fully assembled vector was then transformed into *A. tumefaciens* GV3101 pSOUP+. These were transformed into *A. thaliana* Col-0 wild type plants via floral dip as described in Davis et al. (2009). The resulting seeds were selected as described in Gross et al. (2018). Plants carrying *proCRC:GUS* were crossed with *A. thaliana* Col-0 loss-of-function mutants of putative CRC regulators. Young inflorescences of genotyped F2 plants were harvested in ice cold 90% acetone and incubated for 20 min at room temperature. The GUS staining was performed according to Weigel und Glazebrook (2002). After the staining, the inflorescences were embedded in paraplast for sectioning according to Weigel und Glazebrook (2002). 10 µm thick sections from the embedded tissues were analyzed with a Leica microscope DCM5500.

## Expression analysis

RNA *in situ* hybridization to detect the CRC mRNA in carpel tissue was performed as described in Brewer et al. (2006) with modifications (see Suppl. Figure 4). Probes were generated using T7 RNA Polymerase (for sequences see Suppl. Table 2).

*CRC* expression levels were analyzed via qRT-PCR in mutants of *CRC* regulators. Total RNA from buds of wild type, *arf8, agf2, athb16, bbx19, cal, cil1, ett, ful, haf, hat4, idd12, ino, jag, nf-y9, nga2, rve4, tmo5, ult1*, and *yab5* plants was isolated in quadruplicates using the NucleoSpin RNA Plant kit (Macherey-Nagel GmbH & Co., KG, Düren, Germany) and transcribed into cDNA using RevertAid H Minus Reverse Transcriptase (Thermo Fisher Scientific Inc., Schwerte, Germany) with random hexamer primer. A 1:50 cDNA dilution was added to the Luna master mix (NEB Inc., Frankfurt am Main, Germany) and the qRT-PCR was run on a Lightcycler 480 II (Roche Diagnostics Deutschland GmbH, Mannheim, Germany). ACTIN2 was used as reference gene. Primer efficiencies were determined for CRC 2.1 and ACT2 2.1. Primer sequences are listed in Supplemental table 2. The raw data was analyzed using the Pfaffl method (Pfaffl 2001) and according to Taylor et al. (2010) (for detailed description see Suppl. Fig. 6).

### *In silico* analysis of genomic loci, GO enrichment and co-expression analysis

The genomic loci of *APETALA2, FLOWERING LOCUS T, FLOWERING LOCUS C*, and *CRC* were screened for the presence of miRNA binding sites using psRNA Target (Dai et al. 2018). DNA methylation patterns were analyzed using data from the 1001 Arabidopsis Methylomes Project (Kawakatsu et al. 2016). Histone modifications were identified with PlantPAN3.0 (Chow et al. 2019).

For functional categorization, the putative regulators were imported into Panther (Mi et al. 2013; Thomas et al. 2003) and Gene Ontology terms were attributed from the GO biological process annotation data. Fisher’s Exact test was used and the Bonferroni correction for multiple testing with P < 0.05 was applied.

*CRC* co-expressed genes were identified via Pearson correlation using stage specific carpel (stage 5, 9, 11, and 12) RNA-seq data (Kivivirta et al. 2019) and SAM,leaf, inflorescence, young flowers, and mature flowers RNA-seq data (Klepikova et al. 2016). Genes with positive correlation between 1 – 0.8 and with negative correlation between −0.8 – −1 were used for further analyses. Co-expressed genes present in the Y1H dataset were further analyzed in a heatmap generated with Heatmapper (Babicki et al. 2016) using average linking and Pearson distances. The respective genes were scaled per row. BioGRID (Oughtred et al. 2019) was used to identify protein-protein-interactions between the Y1H identified proteins for the assembly ofe co-regulatory networks.

## Contributions

TG performed all experiments, analyzed the data, and wrote the manuscript, AB devised and designed the work, analyzed the data, and revised the manuscript.

## Acknowledgements

We thank Paula Elomaa and Suvi Broholm for introducing TG to the Y1H technique and Martin Yanofsky and Birgit Poppenberger for suppling us with seeds of the *hbb* line, and Daniel Schubert for the *cal* line, and Jan Lohmann for the Greengate vectors. We thank Agnieszka Golicz for helpful comments on the manuscript. Furthermore, we thank Denise Herbert and Kimmo Kivivirta for supplying us with the RNAseq data. Moreover, we thank Andrea Gómez Felipe for introducing her RNA *in situ* protocol to us. For excellent technical support, we thank Andrea Weisert and Claudia Jung-Blasini. For their assistance in the lab, we thank the students Le Han Nguyen, Larissa Witzel, Dennis Oderwald, and Julian Garrecht.

