## Supplementary figures and images for "Transcription factor action orchestrates the complex expression pattern of *CRABS CLAW*, a gynoecium developmental regulator in Arabidopsis"

### Supplemental Figure 1

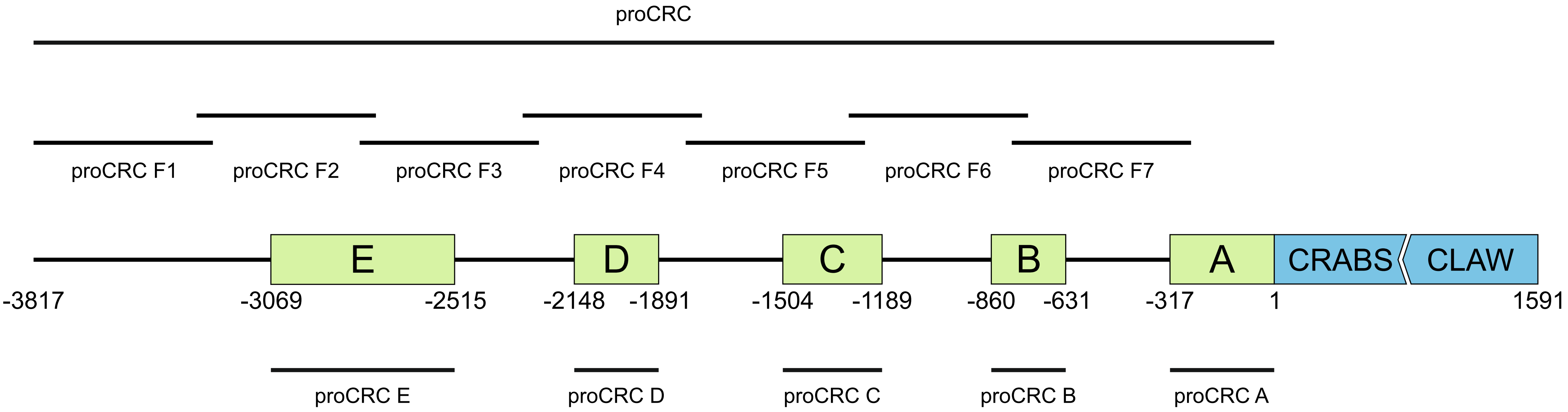

### Supplemental Figure 2

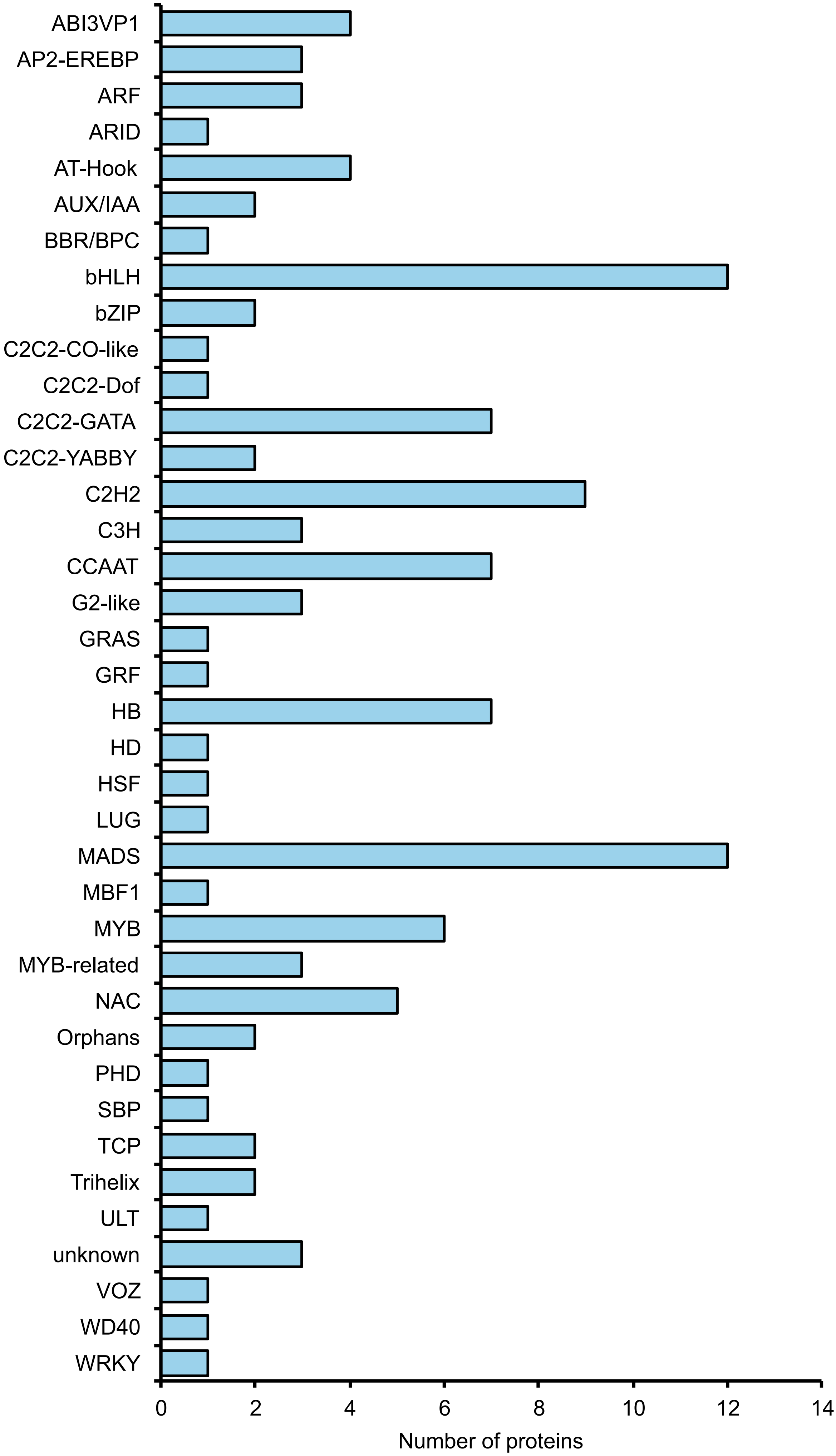

### Supplemental Figure 3

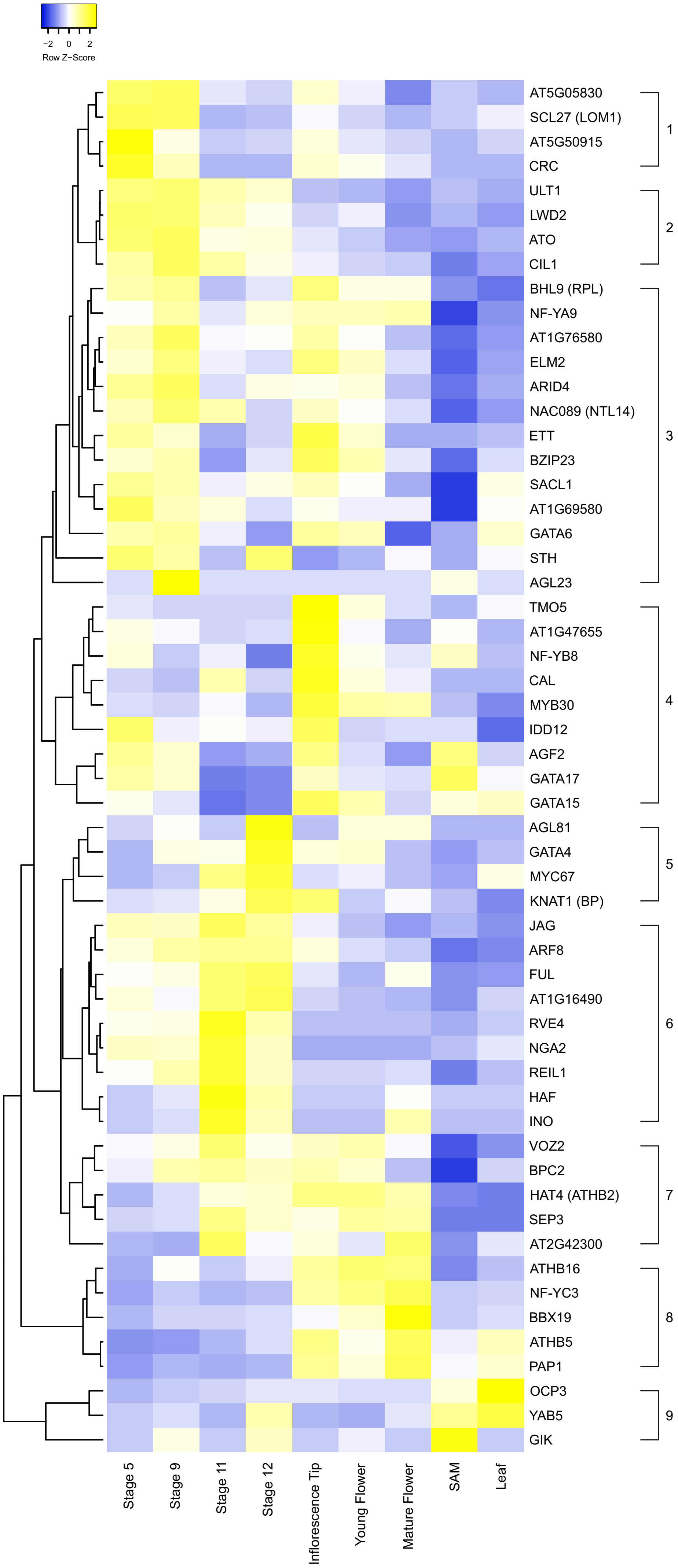

### Supplemental Figure 4

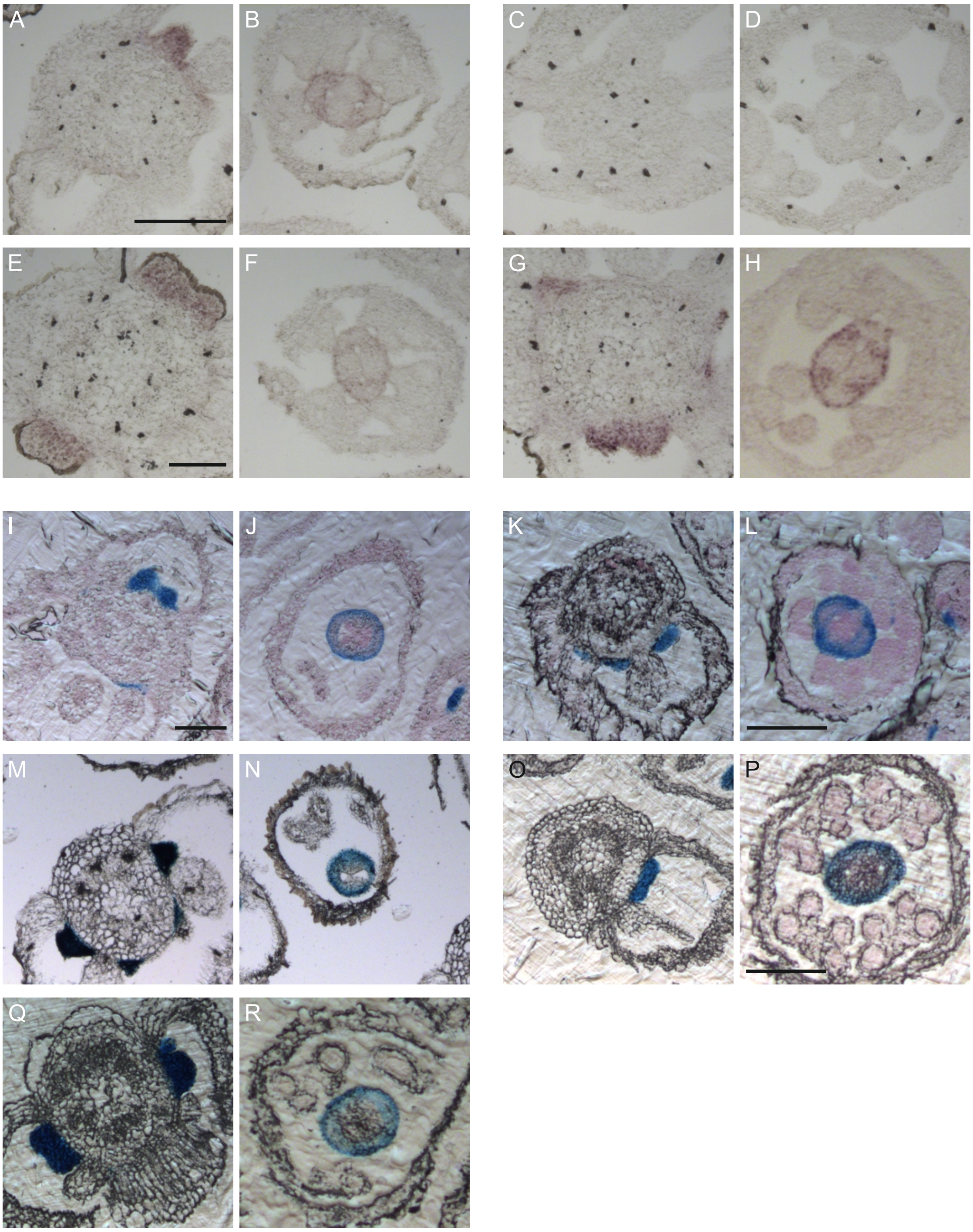

### Supplemental Figure 5

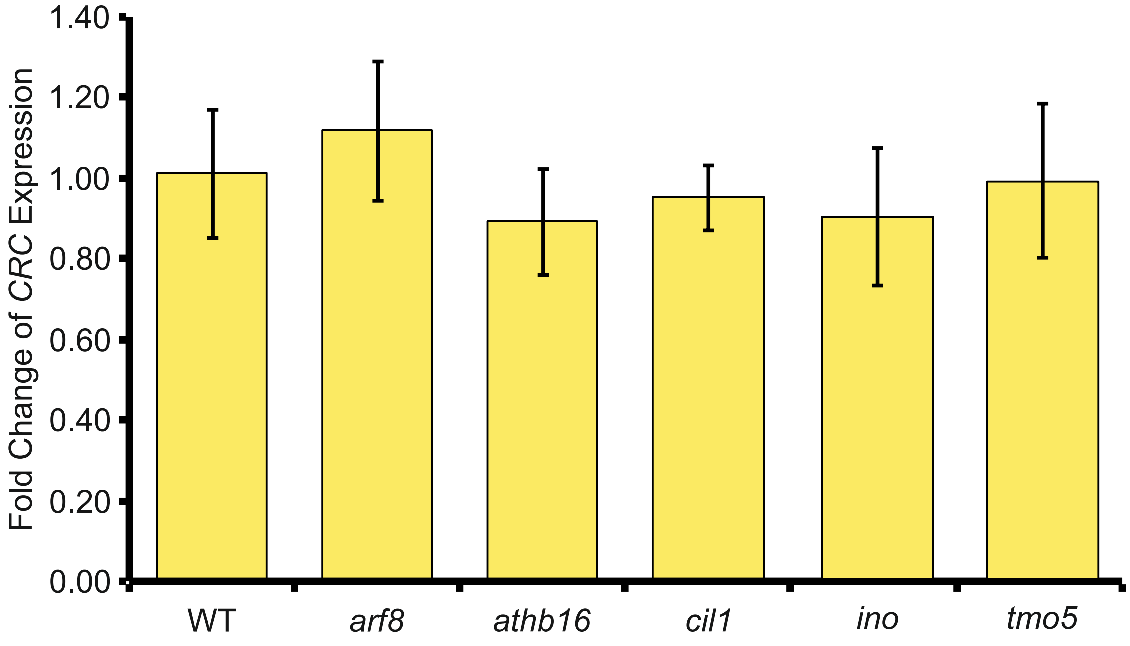
